# Prevalence of GIT nematodes and associated risk factors of exotic chickens in selected farm of poultry in and around Ambo, Ethiopia

**DOI:** 10.1101/2024.07.02.601802

**Authors:** Abraham Belete Temesgen, Zerihun Getie Wassie, Saleamlak Abebe

## Abstract

Poultry are raised worldwide in backyards and commercial systems with fewer social and religious taboos than other livestock. However, the chicken industry faces significant challenges from nematode parasites. A cross-sectional study with a random sampling technique was conducted from April to June 2019 to estimate gastrointestinal (GI) nematode parasites in chickens in selected farms in and around Ambo, Ethiopia. The fecal analysis results revealed that out of 70 samples collected, 60% were positive for gastrointestinal (GI) nematode eggs. Prevalence varied significantly by body condition, with the highest rates in chickens in poor condition (78.57%), followed by medium (54.54%) and good condition (40%). Location also played a significant role, with Ambo University Poultry Farm having the highest prevalence (83.87%), followed by Abebe Private Farm (55%) and Guder Campus Poultry Farm (26.31%). The main nematode species identified were *Ascaridia galli* (57.1%) and *Heterakis gallinarum* (2.9%). Infestation rates differed significantly by sex, age, location, and body condition, with males having higher rates of *Ascaridia galli* (61.53%) than females (56.14%), and *Heterakis gallinarum* exclusively affecting females (3.51%). Adults showed significantly higher rates of *Ascaridia galli* (85.71%) than young chickens (38.09%), with some infestation of *Heterakis gallinarum* (7.14%) observed in adults but absent in young chickens. This prevalence rate suggests limited awareness among chicken producers and insufficient control strategies in the study area. Hence, implementing targeted control strategies is advisable.

## 1. Introduction

Poultry are kept in backyards or commercial production systems in most areas of the world. Compared to a number of other livestock species, fewer social and religious taboos are related to the production, marketing, and consumption of poultry products. For these reasons, poultry products have become one of the most important protein sources for humans throughout the world [11, 20]. In developing countries, poultry production offers an opportunity to feed the fast-growing human population and to provide income resources for poor farmers. Moreover, poultry in many parts of the modern world are considered as the chief source of not only cheaper protein of animal origin but also high-quality human food [7].

Among the important species of livestock kept in Ethiopia, poultry production systems are identified in the country. These include backyard poultry production systems, small-scale, and large-scale intensive production systems [28]. The population of poultry in Ethiopia is estimated to be 44.89 million, excluding the pastoral and agro-pastoral areas. With regard to breed, 96.46%, 0.57%, and 2.97% of the total poultry are reported to be indigenous, hybrid, and exotic, respectively [8]. Despite the presence of a large number of chickens in Ethiopia, the contribution to the national economy or the exploited benefit is very limited due to nutritional limitations and diseases [21].

Among parasitic diseases of poultry, nematode parasites are one of the major problems of the chicken industry worldwide, characterized by riffled feathers, loss of appetite, poor growth, and reduced egg production [1]. Moreover, nematodes (roundworms) are the most important group of helminth parasites of poultry. This is due to the large number of parasitic species that cause damage to the host, especially in severe infections. Most roundworms affect the gastrointestinal tract, with occasional parasites affecting the trachea or eye. Each species of roundworm tends to infect a specific area of the gastrointestinal tract. Different species of the same genus may infect several different areas of the tract. In general, the different species of roundworms have very similar life cycles [5]. Of the helminth parasites of poultry birds, nematodes constitute the most important group of helminth parasites of poultry, both in the number of species and the extent of damage they cause; the main genera include Ascaridia and Heterakis [15].

Generally, nematode infections in poultry are widely distributed in different parts of the world, and numerous research efforts have been undertaken to prevent poultry mortality from parasitic diseases. The prevalence of two nematode genera, Ascaridia and Heterakis, has been extensively studied [2]. Among diseases, internal parasites are known to reduce the productivity of poultry kept under various management systems. Infection by parasites occurs after ingestion of nematode eggs or intermediate hosts such as cockroaches, grasshoppers, ants, and earthworms. Nematode infection results in a reduction in food intake, injury to the intestinal wall, and hemorrhage, leading to poor absorption of nutrients and weight loss [24].

Gastrointestinal parasite infestation is a common problem in poultry, especially when nematode infections occur in high proportions in animals reared in intensive management systems. In Ethiopia, the poultry industry is developing for both local and exotic chickens, but only a few surveys have been carried out to determine the burden of nematode parasites in chickens in this country [17, 19]. Therefore, the objectives of this study were to estimate the prevalence of major gastrointestinal nematode parasites in poultry and to assess the risk factors associated with the incidence of the parasites in the study area.

## 2. Materials and methods

### 2.1. Study area

The study was conducted on selected farms in and around Ambo, Ethiopia (Figure1), from April to June 2019. Ambo is the administrative center of the zone and is located at a latitude and longitude of 8°59′N 37°51′E, with an elevation of 2101 meters above sea level (asl) and 114 km west of Addis Ababa. The Ambo Woreda has 34 administrative kebeles. The agro-ecology of the study area is 23% highland, 60% midland, and 17% lowland. It has an annual rainfall and temperature ranging from 800-1000 mm and 20-29 °C, respectively. The livestock population of the district includes 145,371 cattle, 50,152 sheep, 27,026 goats, 105,794 chickens, 9,088 horses, 2,914 donkeys, and 256 mules [4].

**Figure1:**
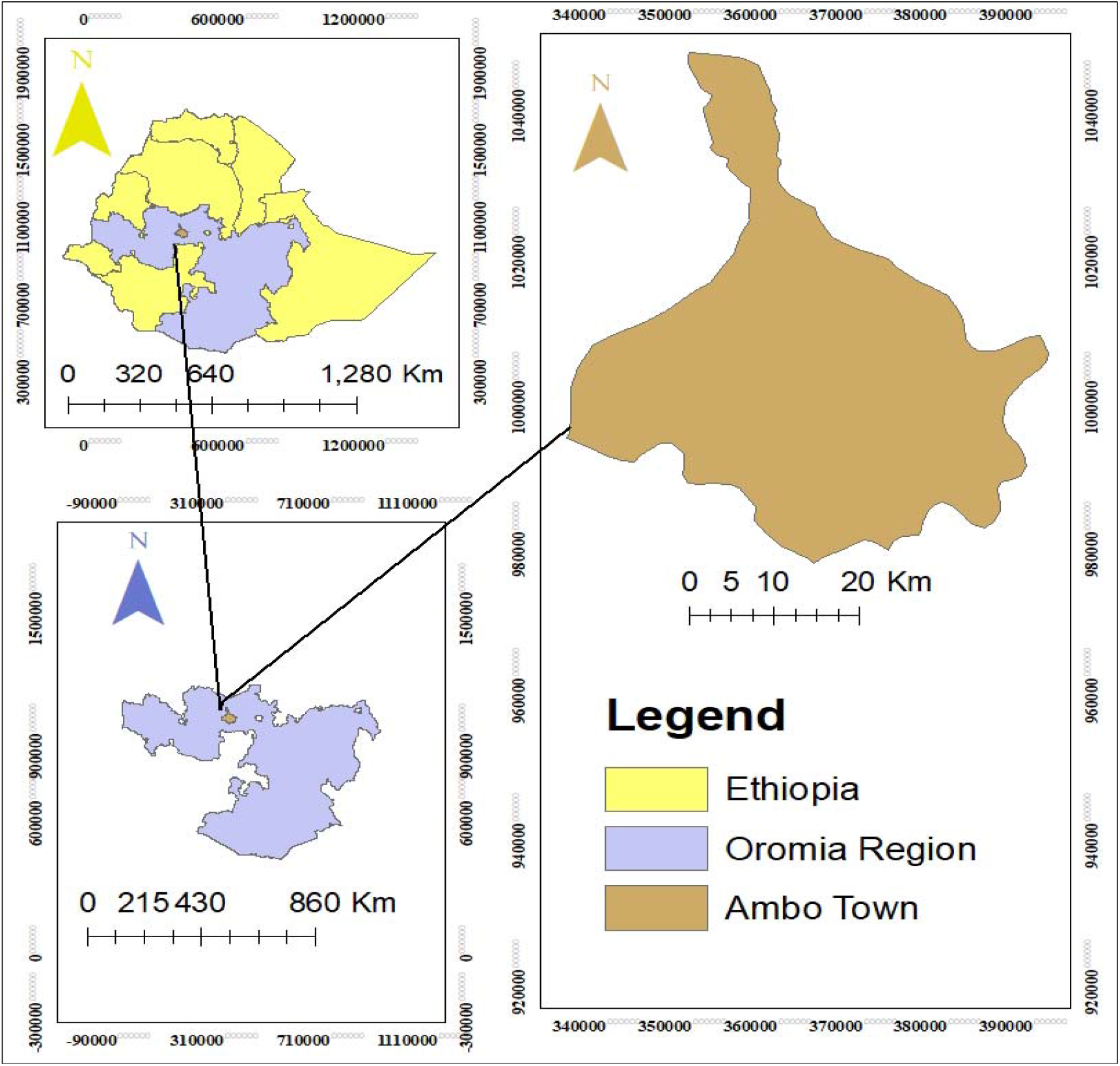
Map of the study area

### 2.2. Study population

The study included all exotic breed chickens of both sexes across various age groups, housed under intensive management on selected farms. Chickens were categorized into two groups: young (under 12 months) and adult (12 months and older). Age determination relied on farm records and subjective assessments such as crown size and spur length [9].

### 2.3. Study design and Sample size determination

A cross-sectional study design was used for this study. Sex, different age groups, body condition, breed, and management systems were recorded as test variables during the data collection of target chickens. A random sampling technique was used to recruit animals for the study. The required sample size for this study was estimated according to the formula of [26], with an expected prevalence of 68.5% from a previous study conducted by [19] in a comparable agro-ecological area, and a desired absolute precision (*d*) of 0.05 at a 95% confidence level.

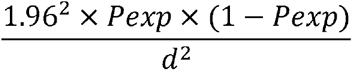

Where *n* represents the required sample size, *P*exp denotes the expected prevalence, and *d* stands for the desired absolute precision. Using this formula, the sample size was calculated to be 332. However, only 70 chickens were selected purposefully due to a shortage of time.

### 2.4. Data collection

After assessing the selected chickens for general body condition and clinical signs suggestive of gastrointestinal nematodes, samples were collected and sent to the Ambo University Department of Veterinary Laboratory Technology for parasitological analysis. Fecal samples were obtained using gloves directly from the vent or top surface of freshly voided feces and placed in sealed screw-cap containers to maintain their integrity. These samples were promptly transported to the Parasitology laboratory at Ambo University’s College of Veterinary Laboratory Technology and stored at 4°C until examination. During collection, detailed information regarding risk factors such as body condition, sex, age, and management practices was meticulously recorded for each sample. The samples underwent processing using the floatation technique, following the procedure outlined by [18].

### 2.5. Data management and analysis

The raw data gathered for the study were entered into a Microsoft Excel database and organized using the Excel spreadsheet program. Subsequently, the data were imported into STATA Version 18.0 for analysis. A chi-square (χ2) test was employed to evaluate the statistical association between infection rates and various factors. A significance level of p < 0.05 at a 95% confidence interval was used to determine statistical significance between variables.

## 3. Results

The fecal analysis results revealed that out of 70 samples collected, 42 (60%) were positive for gastrointestinal (GI) nematode eggs. The prevalence of nematode infection varied significantly with body condition: chickens in poor body condition had the highest prevalence at 78.57%, followed by those in medium condition at 54.54%, and those in good condition at 40%. This difference was statistically significant. Location also played a significant role in the prevalence of GI nematodes. The highest prevalence was observed at Ambo University Poultry Farm (83.87%), compared to Abebe Private Farm (55%) and Guder Campus Poultry Farm (26.31%), with the differences being statistically significant. Among the risk factors evaluated, age, body condition, and location showed statistically significant differences in prevalence, whereas sex did not exhibit a significant difference (p>0.05) (Table 1 and Figure 2).

**Table 1:** The prevalence gastro-intestinal nematode infection by Risk Factors.

| Factors | Category | Examined | Positive | $\chi^2$ | P-value |
| --- | --- | --- | --- | --- | --- |
| **Sex** | Female | 57 | 34 (59.64%) | 0.016 | 0.900 |
|  | Male | 13 | 8 (61.53%) |  |  |
| **Age** | Young | 42 | 16 (38.09%) | 20.992 | 0.000 |
|  | Adult | 28 | 26 (92.85%) |  |  |
|  | Abebe private | 20 | 11 (55%) |  |  |
| **Place (Farm)** | Ambo university | 31 | 26 (83.87%) | 16.55 | 0.000 |
|  | Guder campus | 19 | 5 (26.31%) |  |  |
| **Body condition** | Poor | 28 | 22 (78.57%) | 7.630 | 0.022 |
|  | Medium | 22 | 12 (54.54%) |  |  |
|  | Good | 20 | 8 (40%) |  |  |
| **Total** |  | **70** | **42(60.0%)** |  |  |

**Figure 2:**
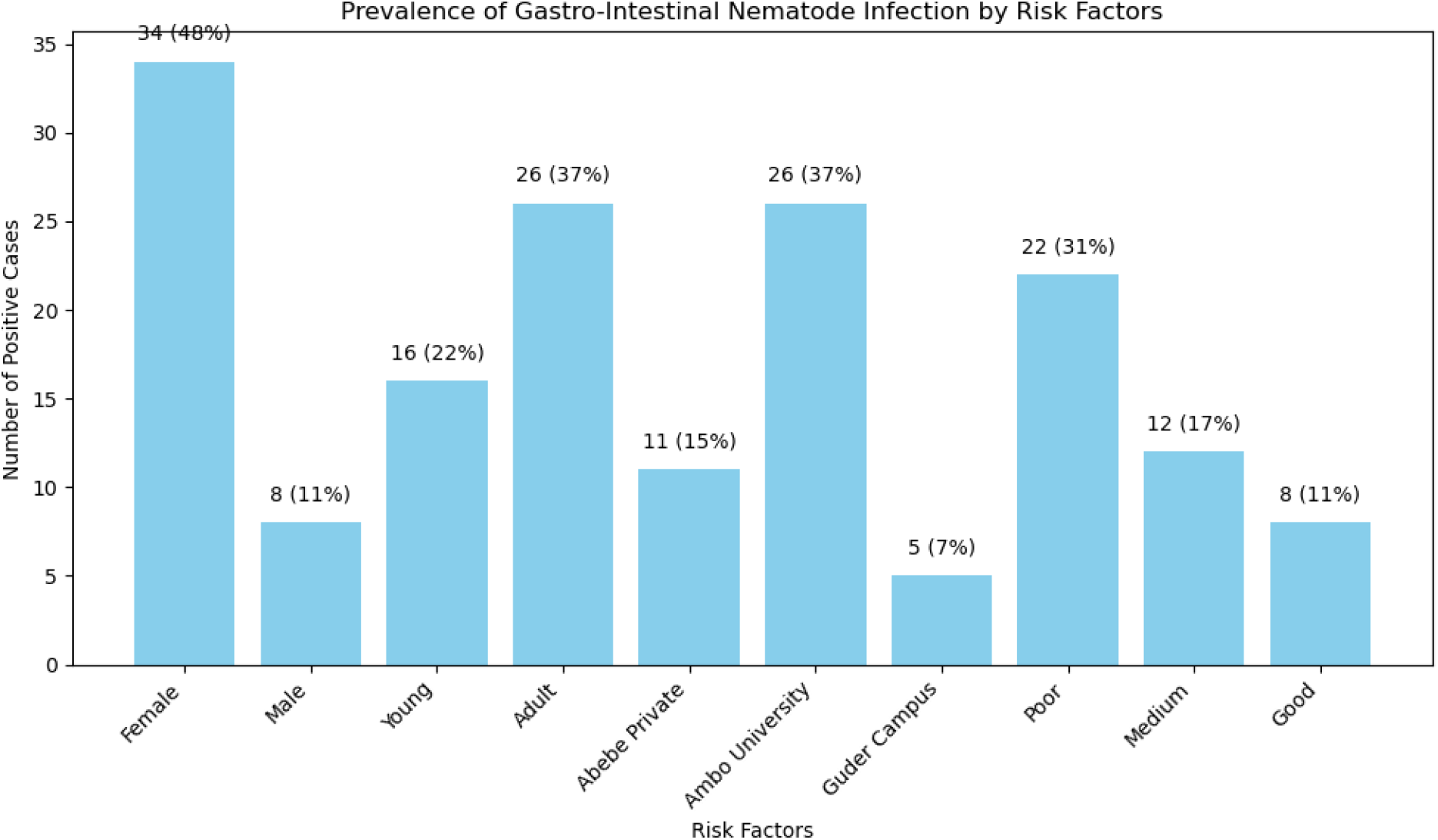
The Prevalence of Nematode species infestation by Risk Factors

The overall prevalence of two nematode species identified from fecal examinations in chickens were *Ascaridia galli* (57.1%) and *Heterakis gallinarum* (2.9%). The infestation rates of these nematodes vary significantly based on sex, age, location, and body condition. Males have a slightly higher infestation rate of *Ascaridia galli* (61.53%) compared to females (56.14%), while *Heterakis gallinarum* is only observed in females (3.51%). Age-wise, adults exhibit a significantly higher infestation rate of *Ascaridia galli* (85.71%) compared to young chickens (38.09%) and also show some infestation of *Heterakis gallinarum* (7.14%), which is absent in young chickens. Location-wise, chickens from Ambo University have the highest infestation rate of *Ascaridia galli* (77.41%) and a noticeable infestation of *Heterakis gallinarum* (6.45%), whereas those from Guder Campus have the lowest rates for both parasites (26.31% and 0%, respectively). Chickens in poor body condition are the most affected, with the highest infestation rates for *Ascaridia galli* (71.42%) and *Heterakis gallinarum* (7.14%), while those in good condition have the lowest rates for both nematodes (Table 2 and Figure 3).

**Table 2:**
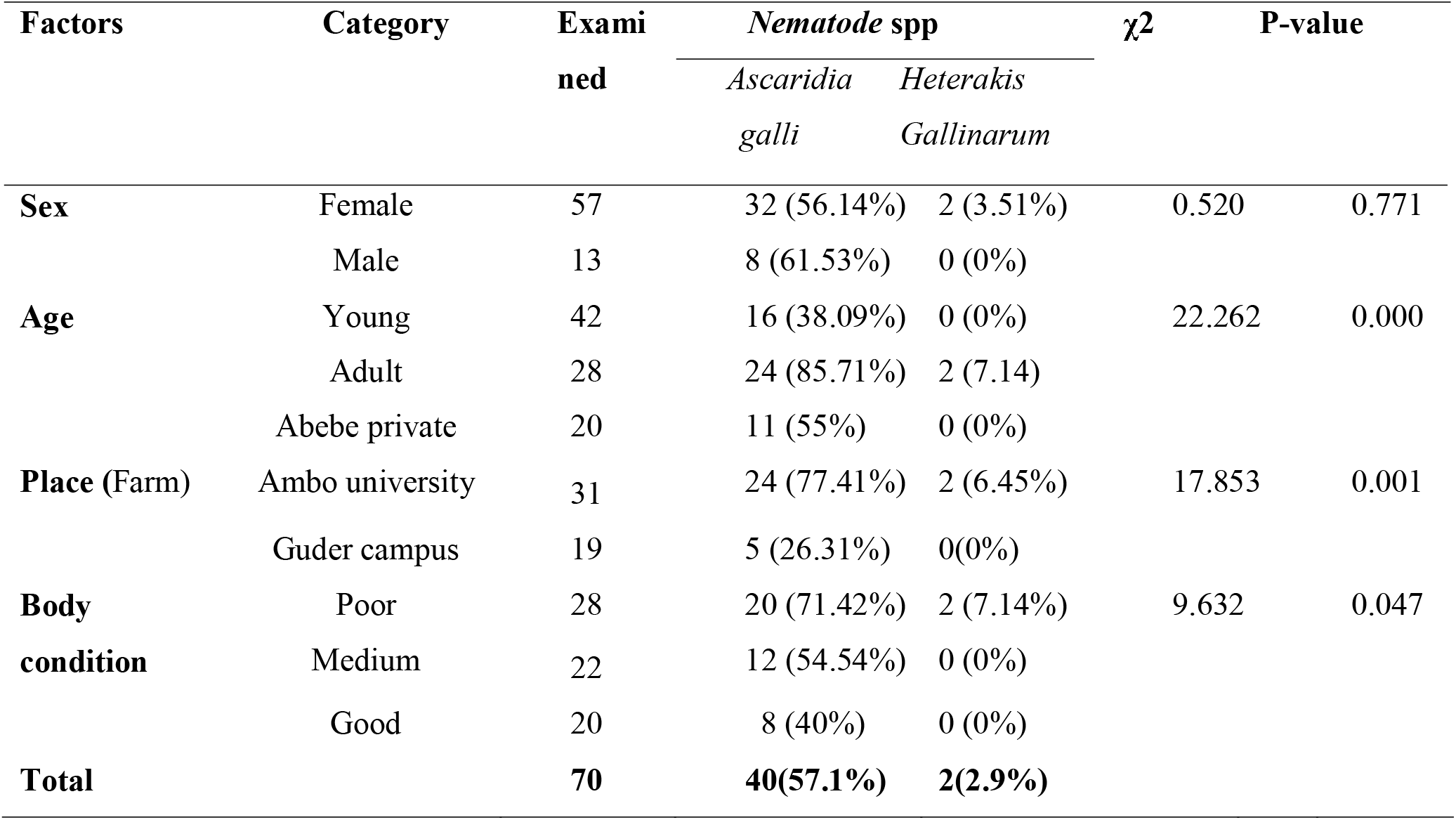
The Prevalence of Nematode species infestation by Risk Factors.

| Factors | Category | Examined | Nematode spp | | $\chi^2$ | P-value |
| --- | --- | --- | --- | --- | --- | --- |
|  |  |  | *Ascaridia galli* | *Heterakis Gallinarum* |  |  |
| **Sex** | Female | 57 | 32 (56.14%) | 2 (3.51%) | 0.520 | 0.771 |
|  | Male | 13 | 8 (61.53%) | 0 (0%) |  |  |
| **Age** | Young | 42 | 16 (38.09%) | 0 (0%) | 22.262 | 0.000 |
|  | Adult | 28 | 24 (85.71%) | 2 (7.14%) |  |  |
|  | Abebe private | 20 | 11 (55%) | 0 (0%) |  |  |
| **Place (Farm)** | Ambo university | 31 | 24 (77.41%) | 2 (6.45%) | 17.853 | 0.001 |
|  | Guder campus | 19 | 5 (26.31%) | 0(0%) |  |  |
| **Body condition** | Poor | 28 | 20 (71.42%) | 2 (7.14%) | 9.632 | 0.047 |
|  | Medium | 22 | 12 (54.54%) | 0 (0%) |  |  |
|  | Good | 20 | 8 (40%) | 0 (0%) |  |  |
| **Total** |  | **70** | **40(57.1%)** | **2(2.9%)** |  |  |

**Figure 3:**
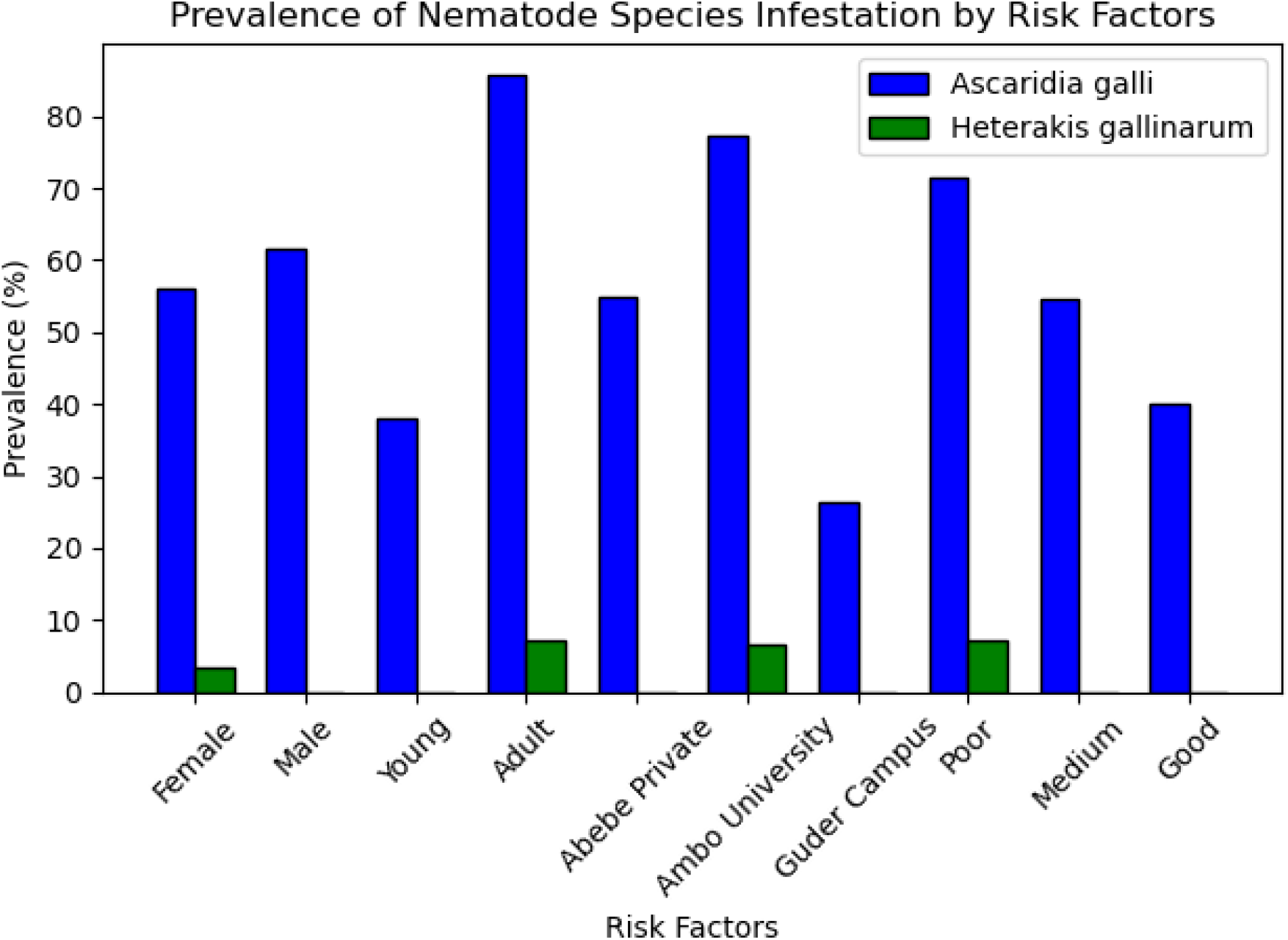
The prevalence of nematode species infestation by Risk Factors

## 4. Discussion

In this cross-sectional study, the overall prevalence of infection with gastrointestinal (GI) nematode eggs was 60.0%. This finding was higher than previous reports by [15], who reported a prevalence of 59.64% and 53% in Ethiopia and Nigeria, respectively. Nonetheless, this finding was lower than the report of [14], which reported 61.9%, and [27], who reported 64.7% in Nigeria and in the Oromia region of Ethiopia, respectively. Additionally, the current study was slightly lower than the result recently reported by [19] of 68.5% from the same area in Ambo, West Shoa Zone, Oromia Regional State, Ethiopia. This discrepancy could be related to differences in the management system, study method, sample size, and control practices in the area.

In the present study, females seemed to have a higher prevalence (59.64%) than males (61.53%), which could be related to the higher susceptibility of female animals. However, there was no significant difference (χ2 = 0.016 and P > 0.05) in the prevalence of gastrointestinal nematode parasites between sexes. This finding aligns with [15, 23], who reported that female chickens were more infected with GI nematode parasites than males. Female chickens are known to be more voracious in their feeding habits, especially during egg production, compared to males. However, this study contrasts with another report from Haromaya by [25], found higher GI nematode infection in males (52.1%) than females (39.9%). This difference may be due to sample size and nutritional deficiency. Another report by [12] indicated no usual natural affinity of GI nematode species to either sex of the host chickens.

Among the age groups, adult chickens had a higher prevalence (92.85%) than young age groups (38.09%), with a statistically significant difference (p < 0.05). This could be due to adult chickens being exposed to infective larval stages for a longer time, contributing to a higher prevalence in older age groups. This finding is consistent with [22] but contrary to [18], who reported a higher level of GI nematode prevalence in young chickens. This could be due to young chickens having a lower level of immunity compared to adults. Similarly, [16] observed a higher prevalence of GI nematodes in young chickens than in adults in Kenya.

The most prevalent nematode species encountered in the present study was *Ascaridia galli* (57.1%). The prevalence of *Ascaridia galli* was higher than the previously reported works in central Ethiopia by [3] (55.26%) and [25] from Haromaya (38.0%). This might be due to differences in management systems, deworming practices, and/or agro-ecological conditions of the study area. The high occurrence of the parasites in the study area may be related to the wet season when the survey was conducted. The prevalence of *H. gallinarum* (2.9%) in this study was lower than in another study in Ethiopia [10] (4.3%) and higher than another study in Kenya [13] (1.43%). However, it is lower than the 51.6% reported by [6] in Ethiopia, possibly due to agro-ecological variation.

## 5. Conclusion

The current study indicated that the overall prevalence of gastrointestinal nematode (GIT) infections in chickens in the study area was 60.0%. GIT nematode infections are found to be an important problem in the study area. The results of this study showed that *Ascaridia galli* was the most dominant species. The compared risk factors in the current study, such as body condition, place, and age, showed significant variation with nematode infection. This prevalence rate suggests limited awareness among chicken producers and insufficient control strategies in the study area. Therefore, implementing targeted control strategies is advisable.

## Data Sharing Statement

All data generated or analyzed during this study are available upon request from the corresponding author.

## Ethics Statement

Not applicable.

## Acknowledgement

The author expresses heartfelt gratitude to all co-authors of this manuscript for their invaluable contributions and support.

## Authors’ information

Abraham Belete Temesgen (ABT); Zerihun Getie Wassie (ZGW); Saleamlak Abebe (SA)

## Author Contributions

ABT contributed to writing the original draft, resources, methodology, investigation, conceptualization, formal analysis, supervision, and data curation. ZGW contributed to supervision, writing the original draft, resources, methodology, investigation, and conceptualization. SA contributed to writing the original draft, resources, methodology, investigation, and conceptualization.

### Funding

None

### Disclosure

The author declares no conflicts of interest in this work.

